# A method for non-invasive prenatal diagnosis of monogenic autosomal recessive disorders

**DOI:** 10.1101/635342

**Authors:** Anthony Cutts, Dimitrios V. Vavoulis, Mary Petrou, Frances Smith, Barnaby Clark, Shirley Henderson, Anna Schuh

## Abstract

Non-invasive prenatal testing (NIPT) to date is used in the clinic primarily to detect foetal aneuploidy. Few studies so far have focused on the detection of monogenic autosomal recessive disorders where mother and foetus carry the same mutation. In particular, NIPT is currently not available for the detection of Sickle Cell Anaemia (SCA), the most common monogenic disorder world-wide and the most common indication for invasive prenatal testing in high-income countries. Here, we report the clinical validation of a novel diagnostic approach that combines ultra-sensitive amplicon-based sequencing of cell-free DNA (cfDNA) with internal controls and bias factor correction to calculate the probability for the presence of allelic imbalance from maternal plasma without prior knowledge of the paternal genotype. Identification of the foetal genotype was determined using a hierarchical probabilistic model based on the relative number of reads from the sequencing, along with the foetal fraction. NIPT was performed on a cohort of 57 patients, all of whom had previously undergone invasive prenatal testing so that the foetal genotype was known. Overall, NIPT demonstrated 100% sensitivity and negative predictive value for foetal fractions higher than 0.5%, and 100% specificity and positive predictive value for foetal fractions higher than or equal to 4%. Our methodology can be used as a safe, non-invasive screening tool in any clinical scenarios where early prenatal diagnosis of SCA or other recessive disorders is important.

## Main

Since the discovery of circulating cell-free DNA (cfDNA) in the plasma of pregnant women in the late 1990s^1,2^, its potential for prenatal diagnosis has been the focus of intensive technological innovation. Screening for chromosomal abnormalities is now introduced in several countries including the UK^3^. However, the so-called ‘combined test’ (which utilises ultrasound scanning to measure foetal nuchal translucency, maternal age and blood tests to measure pregnancy-associated plasma protein A and free beta-human chorionic gonadotrophin) remains the first-stage test. Women at high-risk of carrying an affected baby are offered NIPT and, if this returns a positive result, they are given the option of confirmatory invasive testing by amniocentesis.

Technologies detecting dominant de-novo mutations or dominant mutations in the father from cfDNA have been previously described^4^. For recessive disorders characterised by compound heterozygosity (e.g. cystic fibrosis), the same technology can be applied if the mutations in both parents are different. Where mutations are the same, methods that utilise the linkage of disease-causing mutations to paternal SNPs have been proposed^5,6^. However, this approach requires testing a previously born child for linkage to the paternal SNPs and it is subject to errors due to recombination events. An alternative approach avoiding the need for complex family work-up, would be to measure small allelic imbalances caused by foetal DNA in the maternal circulation. However, this requires precise allele quantification and, so far, rapid NIPT from cfDNA for conditions such as SCA, where the mother and father carry the same mutation, remains elusive.

SCA is an autosomal recessive disease characterised by a single base-pair substitution in the beta globin gene. Due to the protective effect of the mutation against malaria, carrier frequencies in sub-Saharan Africa are 20% or higher. Over 224,200 infants are born annually with SCA worldwide^7^, including at least 1,000 in the US, making SCA by far the most common monogenic disease indication for invasive prenatal testing (IPT) in high-income countries. Invasive prenatal diagnosis by amniocentesis or chorionic villus sampling is costly and carries a 1-2% risk of miscarriage. Only one study so far has reported the use of cfDNA for the detection of allelic imbalance in SCA using digital droplet-PCR^8^ and, to our knowledge, no clinical services currently offer the test worldwide.

Here, we describe the development and validation of a highly sensitive and specific next-generation sequencing (NGS) approach for the non-invasive prenatal diagnosis (NIPD) of SCA. Our methodology does not require the paternal genotype and combines optimised PCR of the affected locus (including sequencing and PCR error correction) with precise estimation of the foetal cell fraction and different internal controls (Figure 1). The raw assay data are analysed using a bespoke statistical methodology (Figure 2), which estimates the respective foetal disease status. We calibrated and, subsequently, tested our method on 57 patients, achieving very high sensitivity (94%) and specificity (88%) (Figure 3 and Table 1). The method is also applicable to other autosomal recessive and autosomal dominant diseases.

**Table 1.** Training and test cohorts used in this study and predictions from application of the NIPT. False positives are indicated in yellow. The false negative is indicated in red.

| Training cohort |  |  |  |  |  |  |
| --- | --- | --- | --- | --- | --- | --- |
| Sample | Foetal fraction (%) | Maternal genotype | True foetal genotype | True disease state | E[P] | Predicted disease state |
| 1 | 9.7 | AS | AA | not SS | 0.18 | not SS |
| 2 | 5.3 | AS | AS | not SS | 0.53 | not SS |
| 3 | 4.8 | AS | SS | SS | 0.79 | SS |
| 4 | 4.1 | AS | SS | SS | 0.86 | SS |
| 5 | 4.0 | AS | SS | SS | 0.82 | SS |
| 6 | 4.0 | AS | SS | SS | 0.81 | SS |
| 7 | 3.8 | AS | AA | not SS | 0.72 | SS |
| 8 | 3.4 | AS | AS | not SS | 0.81 | SS |
| 9 | 2.9 | AS | AS | not SS | 0.60 | not SS |
| 10 | 2.9 | AS | AS | not SS | 0.15 | not SS |
| 11 | 2.6 | AS | AS | not SS | 0.78 | SS |
| 12 | 2.5 | AS | AS | not SS | 0.59 | not SS |
| 13 | 2.2 | AS | SS | SS | 0.82 | SS |
| 14 | 2.1 | AS | AA | not SS | 0.48 | not SS |
| 15 | 2.0 | AS | AA | not SS | 0.15 | not SS |
| 16 | 1.9 | AS | SS | SS | 0.86 | SS |
| 17 | 1.7 | AS | AS | not SS | 0.38 | not SS |
| 18 | 1.7 | AS | AS | not SS | 0.39 | not SS |
| 19 | 1.7 | AS | AA | not SS | 0.14 | not SS |
| 20 | 1.6 | AS | AS | not SS | 0.16 | not SS |
| 21 | 1.5 | AS | AS | not SS | 0.15 | not SS |
| 22 | 1.4 | AS | AS | not SS | 0.59 | not SS |
| 23 | 1.4 | AS | AA | not SS | 0.45 | not SS |
| 24 | 1.3 | AS | AA | not SS | 0.26 | not SS |
| 25 | 0.9 | AS | AA | not SS | 0.62 | not SS |
| 26 | 0.9 | AS | AS | not SS | 0.62 | not SS |
| 27 | 0.9 | AS | SS | SS | 0.86 | SS |
| 28 | 0.8 | AS | SS | SS | 0.66 | SS |
| 29 | 0.5 | AS | SS | SS | 0.16 | not SS |
| Test cohort |  |  |  |  |  |  |
| Sample | Foetal fraction (%) | Maternal genotype | True foetal genotype | True disease state | E[P] | Predicted disease state |
| 1 | 6.7 | AS | SS | SS | 0.72 | SS |
| 2 | 5.6 | AS | AA | not SS | 0.15 | not SS |
| 3 | 5.5 | AS | AS | not SS | 0.60 | not SS |
| 4 | 4.9 | AS | AA | not SS | 0.51 | not SS |
| 5 | 4.9 | AS | AA | not SS | 0.15 | not SS |
| 6 | 4.7 | SS | AS | not SS | 0.11 | not SS |
| 7 | 3.7 | AS | AA | not SS | 0.15 | not SS |
| 8 | 3.2 | AS | SS | SS | 0.79 | SS |
| 9 | 3.1 | AS | AS | not SS | 0.75 | SS |
| 10 | 3.1 | AS | AA | not SS | 0.18 | not SS |
| 11 | 2.9 | AS | AS | not SS | 0.77 | SS |
| 12 | 2.9 | AS | SS | SS | 0.86 | SS |
| 13 | 2.7 | AS | AA | not SS | 0.14 | not SS |
| 14 | 2.6 | AS | AA | not SS | 0.15 | not SS |
| 15 | 2.6 | AS | AS | not SS | 0.23 | not SS |
| 16 | 2.2 | AS | AS | not SS | 0.41 | not SS |
| 17 | 2.2 | AS | SS | SS | 0.86 | SS |
| 18 | 1.9 | AS | SS | SS | 0.81 | SS |
| 19 | 1.9 | AS | AS | not SS | 0.17 | not SS |
| 20 | 1.7 | AS | AS | not SS | 0.27 | not SS |
| 21 | 1.5 | AS | AS | not SS | 0.54 | not SS |
| 22 | 1.4 | AS | SS | SS | 0.70 | SS |
| 23 | 1.2 | AS | AS | not SS | 0.34 | not SS |
| 24 | 1.0 | AS | AA | not SS | 0.60 | not SS |
| 25 | 0.9 | AS | AS | not SS | 0.15 | not SS |
| 26 | 0.8 | AS | AA | not SS | 0.16 | not SS |
| 27 | 0.7 | AS | AS | not SS | 0.15 | not SS |
| 28 | 0.5 | AS | SS | SS | 0.64 | SS |

**Figure 1:**
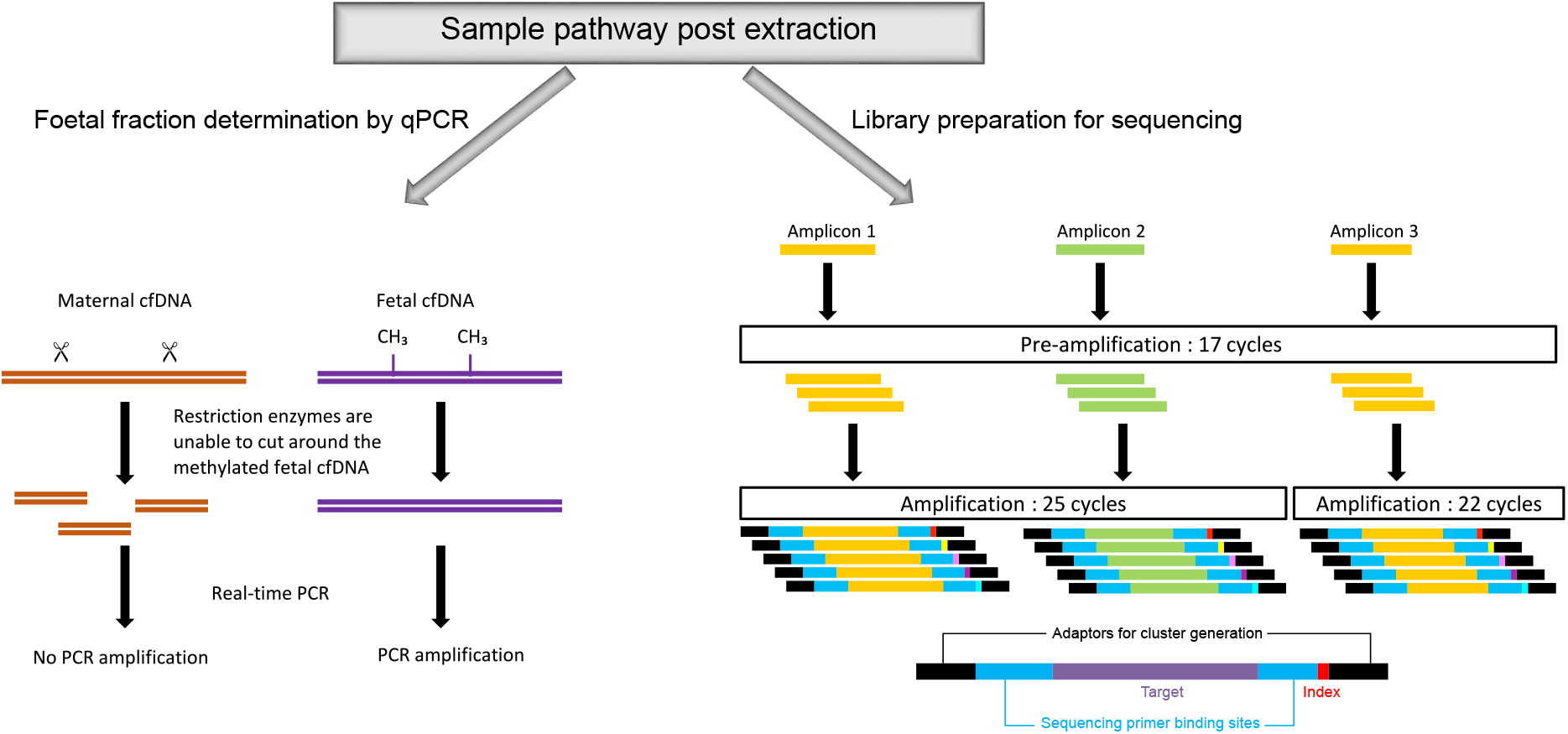
Overview of the next-generation sequencing. Foetal fraction determination was based on methylation status described previously^9^. Library preparation was performed in triplicate, with an initial PCR using standard primers, followed by a second amplification using longer primers containing adapters for cluster generation and indexes so that different patient samples could be multiplexed.

**Figure 2:**
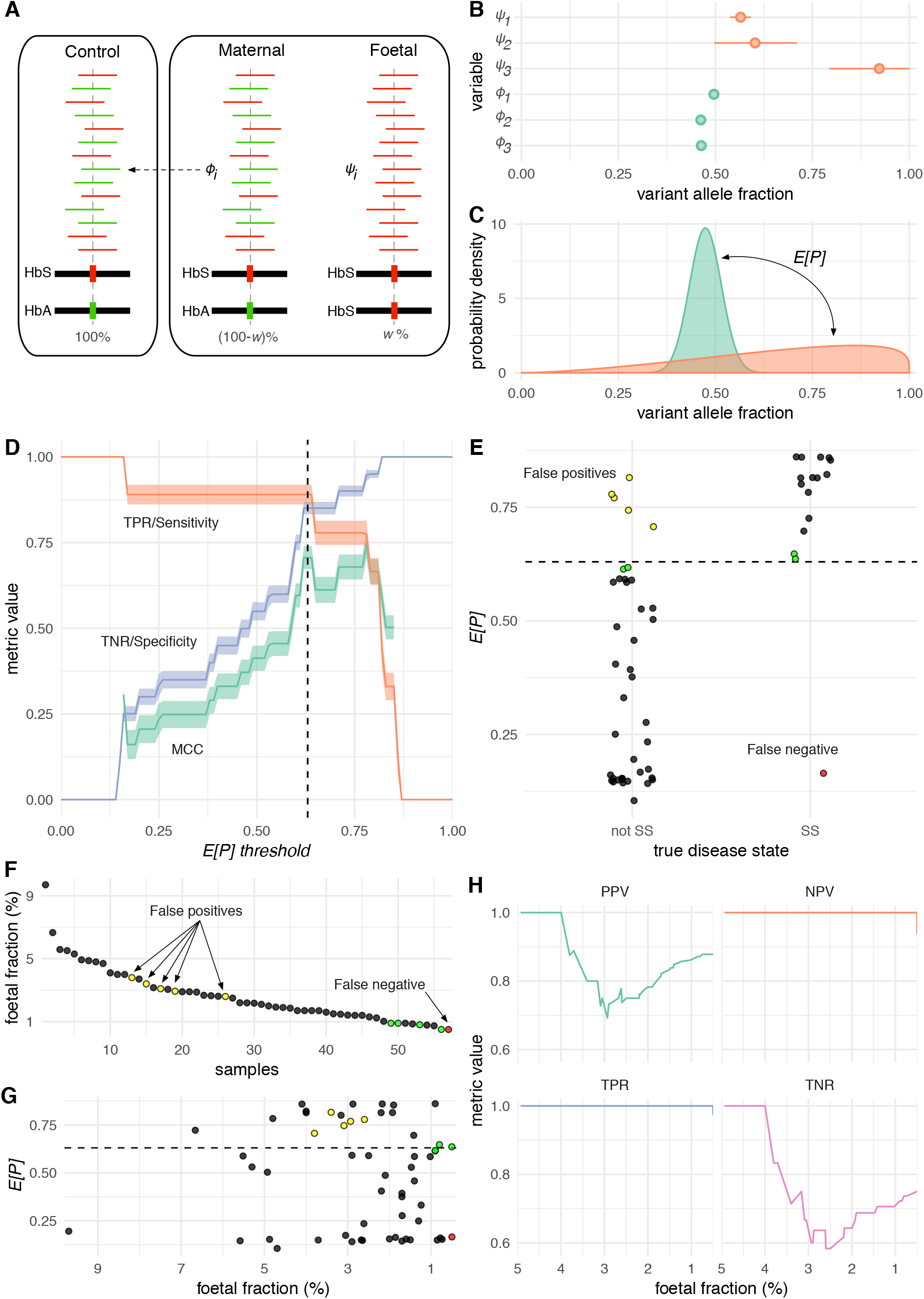
Overview of the statistical analysis. A) The total cfDNA is modelled as a mixture of maternal *ϕ*_*i*_ and foetal *ψ*_*i*_ components with proportions determined by the foetal fraction *w*. B) Estimated maternal and foetal expected fractions of mutated reads for each amplicon in a random subject. C) The estimated probability density functions summarising the expected maternal and foetal fractions in B. The stronger the shift *E*[*P*] of the foetal component to the right of the maternal component, the more likely it is that the foetus has the disease. D) Overview of model training. We identified an optimal *E*[*P*] threshold equal to 0.62. The variance of the various performance metrics was estimated using the bootstrap. E) Overview of applying the calibrated model on both the training and test cohorts. F, G, H) Overview of model performance with decreasing foetal fraction. A false negative arises at a foetal fraction of 0.5%, while false positives arise at foetal fractions less than 4%. Four samples (in green) with *E*[*P*] scores close to the threshold are also characterised by small foetal fractions (≤0.9%)

**Figure 3:**
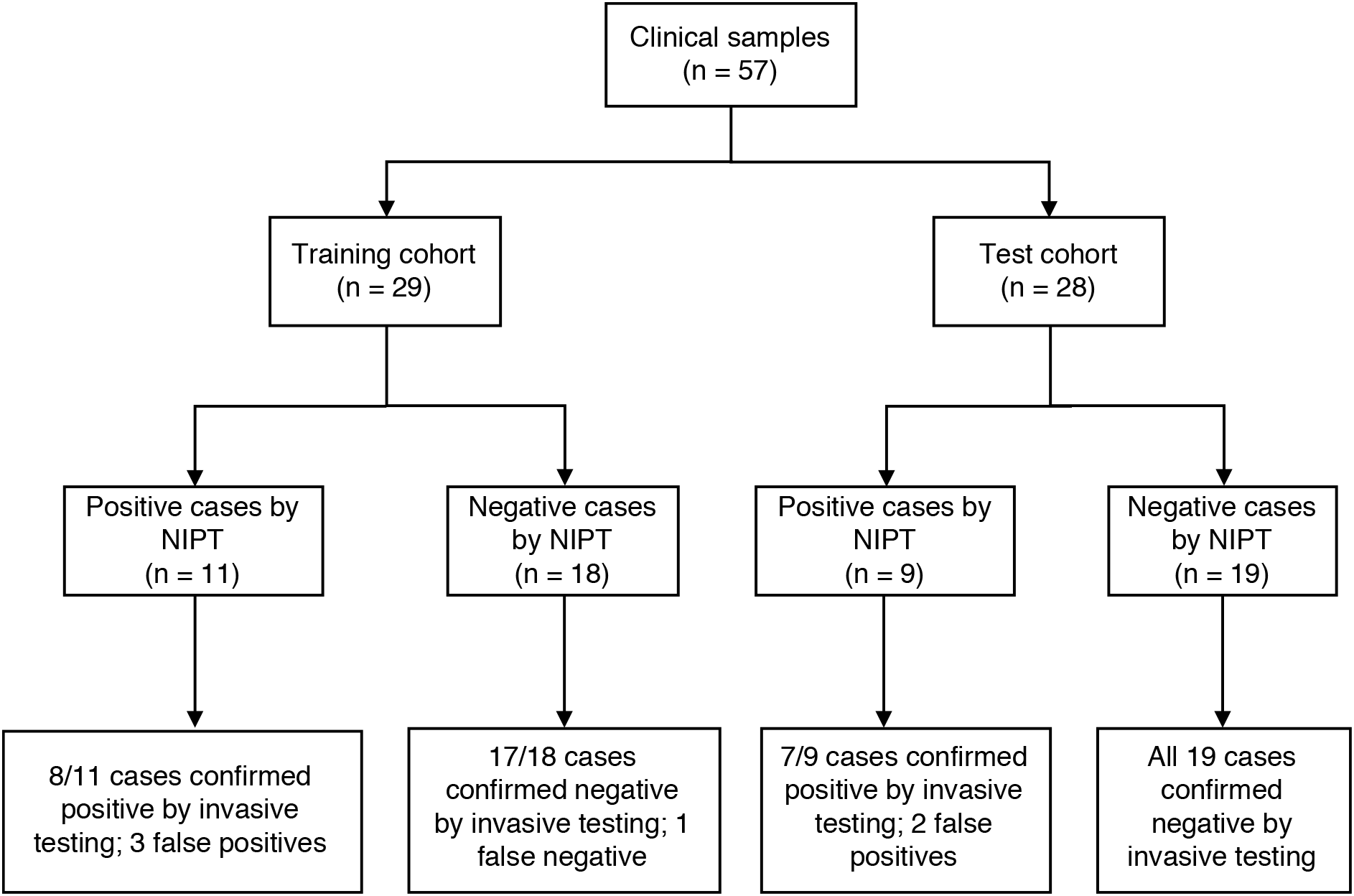
Overview of training and test cohorts used in this study. A total of 57 subjects were recruited, all of which had previously undergone prenatal invasive testing. 29 subjects were used for calibrating the method, i.e. for finding an optimal threshold for the score *E*[*P*] (training phase). The remaining 28 subjects were used for evaluating the predictive capacity of the calibrated model (test phase). Overall, the method returned 5 false positives and a single false negative at foetal fractions ≤0.5%.

Foetal fractions were determined using RT-PCR^9^, followed by library preparation and NGS. Identifying low-level variants using NGS is challenging due to errors during library preparation and sequencing. Additionally, library preparation of highly fragmented cfDNA is more difficult. To overcome these issues, we employed a PCR-based method that generates small amplicons for triplicate analysis to improve noise reduction and error correction. For the library preparation, the first step was a short 17-cycle PCR with short primers specifically designed to preserve the wild-type-to-sickle allele ratio of the template DNA, thus minimizing amplification bias. A second PCR followed with considerably longer primers containing the indices, primer binding sites and adapters required for sequencing. Paired-end ultra-deep sequencing was carried out using the MiSeq 300 v2 chemistry. Levels of the HbS allele were derived from FASTQ files using a Perl script. Up to 8 patient samples were sequenced per run, alongside three controls – with genotypes AA (normal), AS (carrier) and SS (disease) – for normalizing sample data.

NGS is complemented by a bespoke statistical model taking as input the estimated foetal fraction *w*, the number of reads harbouring the HbS allele and the total number of reads covering the locus of the mutation in each amplicon in both the mother and a non-pregnant control with the same genotype as the mother. For each amplicon, we model the expected fraction of reads harbouring the HbS allele in the total cfDNA isolated from the mother as a mixture of foetal (*ψ*_*i*_) and maternal (*ϕ*_*i*_) components in proportions determined by the foetal fraction *w* (Figure 2A). The control shares the same expected fraction of mutated reads *ϕ*_*i*_ with the mother, which helps increase the precision of estimates. For each case, we estimate the foetal (*ψ*_1_,*ψ*_2_,*ψ*_3_) and maternal (*ϕ*_1_,*ϕ*_2_,*ϕ*_3_) expected fractions of mutated reads per amplicon (Figure 2B) and probability density functions summarising each triplet of expected fractions (Figure 2C). For a carrier mother (AS), the density function for the maternal mutated read fractions is concentrated close to 50%, while for a foetus with increased mutated read fractions, the corresponding function is shifted to the right (Figure 2C). The expected magnitude of this shift, *E*[*P*], ranges between 0 and 1 and it predicts the foetus as HbSS (homozygote), if sufficiently high.

In order to determine an optimal threshold value for *E*[*P*], we recruited 29 subjects with known foetal disease status from IPT. We applied NIPT and calculated an *E*[*P*] value for each. Then, for *E*[*P*] threshold values between 0 and 1, we predicted the foetal disease status in each case and calculated the sensitivity (True Positive Rate or TPR), specificity (True Negative Rate or TNR), and Matthews Correlation Coefficient (MCC), a balanced metric for measuring the performance of binary classifiers (Figure 2D). We estimated the variance of all three metrics using the bootstrap (Online Methods). Increasing values of the *E*[*P*] threshold had opposite effect on sensitivity and specificity, reaching an optimal value of 62% at which MCC was maximised (Figure 2D). A second MCC maximum at 77% associated with lower sensitivity (higher false negative rate) was ignored. At the optimal *E*[*P*] threshold, our NIPT correctly identified 8 true positives, 17 true negatives, 1 false negative and 3 false positives (Table 1; Figure 3) achieving 89% sensitivity, 85% specificity, 73% positive predictive value (PPV) and 94% negative predictive value (NPV).

In order to assess the predictive capacity of our NIPT, we recruited 28 additional subjects, which had also undergone IPT, and we applied our method on each using the previously determined optimal *E*[*P*] threshold of 62%. The test returned 7 true positives, 19 true negatives, 2 false positives and no false negatives, achieving 100% sensitivity and NPV, 91% specificity and 78% PPV (Table 1; Figure 3).

On all 57 subjects, the proposed NIPT called 5 false positives and 1 false negative (Figure 2E) corresponding to 94% sensitivity, 88% specificity, 75% PPV and 98% NPV. The single false negative can be attributed to the very low foetal fraction (0.5%) of the corresponding case (Figure 2F). Furthermore, four samples close to the *E*[*P*] threshold (green dots in Figures 2E-G) are also associated with very low (≤0.9%) foetal fractions (Figure 2G). Overall, the NIPT demonstrates 100% sensitivity and NPV for foetal fractions higher than 0.5%, and 100% specificity and PPV for foetal fractions higher than or equal to 4% (Figure 2H).

These results suggest that our methodology can be used as a non-invasive screening tool to exclude the presence of an affected baby. We expect that our test will be implemented similarly to non-invasive aneuploidy diagnosis for confidently excluding an affected pregnancy, thereby considerably reducing the number of IPT performed to confirmatory testing of positive results only. This will particularly also apply to countries with high incidence of sickle cell disease and a demand for prenatal diagnosis (i.e. Nigeria) where access to invasive testing is limited due to high cost and relatively low numbers of trained obstetricians. By comparison, phlebotomy is relatively inexpensive and readily available. This methodology could potentially make NIPT available to a larger number of beneficiaries.

We believe that the future adoption of this NIPT will also have utility in other clinical scenarios:

1. Treatment options for children and young adults who develop early complications from SCA include bone marrow or cord blood transplantation. Several clinical trials using autologous gene-editing of stem cells also reported successful outcomes^10,11^. Knowledge of the foetal genotype before birth would allow advance preparation for umbilical cord blood stem cell sampling for future cellular therapy, without placing the foetus at unnecessary risk through invasive testing.
2. Neonatal screening programmes around the world face logistical challenges because of loss to follow-up of a significant number of babies after birth due to delays in obtaining post-natal screening test results. Knowledge of the foetal genotype prior to birth would allow targeting follow-up efforts to affected pregnancies only.
3. Importantly, our method does not require prior knowledge of the father’s genotype, which will allow its straightforward incorporation into routine antenatal care with a rapid turnaround time.
4. The test could easily be adapted for detection of dominant monogenic disorders, de novo mutations and other autosomal recessive conditions.

## Online Methods

### Ethics statement

Mothers undergoing invasive prenatal diagnosis for SCA were consented for blood sampling in accordance with the Helsinki Declaration for service evaluation.

### Samples

A total of 57 samples were collected from 2012 to 2014, where the gestational ages were between 8 and 17 weeks. Blood was collected in either EDTA or Streck tubes, and processed within 6 or 24 hours of collection, respectively. Blood samples were initially centrifuged at 3000 or 3400 rpm for 10 min at room temperature to separate the plasma. The plasma supernatant was transferred to a fresh tube and micro-centrifuged at 7000 or 14000 rpm for 10 min at room temperature prior to storage at −80°C. cfDNA was extracted using the QIAamp Circulating Nucleic Acid Kit according to the manufacturer’s instructions and eluted into a final volume of 70µl.

### Foetal fraction determination and library preparation

Initially, a Real-Time Polymerase Chain Reaction (RT-PCR) assay was carried out to determine the foetal fraction in the total cfDNA present. This was achieved by assessing the methylation status of the RASSF1A promoter, which is a universal foetal DNA marker, as described previously^9^. For the library preparations, an amplicon-based approach was used where cfDNA samples were amplified by 3 different primer sets in singleplex PCRs. Amplification was carried out in two stages. For the initial PCR, there was 10µl input DNA per amplicon and upon completion of this step, a 2.5µl aliquot was taken and a second PCR performed to attach the appropriate adaptors, primer binding sites and barcodes for sequencing. For both amplifications, primers at a final concentration of 240nmol/L and 0.1 U/µl of Pwo DNA polymerase (Roche) were used with 2X QIAGEN Multiplex PCR Master Mix. The cycling conditions for both PCRs were as follows: 94°C for 3 minutes, then 17 cycles (primary PCR); 25 cycles for amplicons 1 and 2 or 22 cycles for amplicon 3 at 94°C for 45 seconds, 56°C for 45 seconds and 72°C for 1 minute followed by 72°C for 10 minutes (secondary PCR). Following the second PCR, a clean-up was performed and the samples quantified for sequencing using the Qubit Fluorometer. Paired-end ultra-deep sequencing was carried out on a MiSeq, where 84 bases were sequenced from both ends of the DNA fragments.

### Bioinformatics

From the generated FASTQ files, reads above Q30 were processed with a custom Perl script. Using a 2bp amplicon ID tag, the reads were divided into 3 amplicons, based on the following rules: the 84bp sequenced amplicon 1 starts with the bases AC; amplicon 2 ends with the bases TC; and amplicon 3 ends with the bases AC. Using the script, both reads were interrogated to find variants in the DNA fragment at the HbS mutation site (chr11:5,248,232).

### Statistical analysis

Statistical analysis was performed in R^12^ and Stan^13^ using a bespoke statistical model. The model takes as input a) an estimate of the foetal fraction *w* in the total cfDNA, b) the total number of reads *R*_*i*_ covering the HbS locus and the number of reads *r*_*i*_ carrying the HbS allele for each amplicon *i*, as determined from sequencing the total cfDNA extracted from the mother’s blood and c) the total number of reads *S*_*i*_ covering the HbS locus and the number of reads *s*_*i*_ carrying the HbS allele for each amplicon *i*, as determined from sequencing the total cfDNA extracted from the blood of a non-pregnant control subject with the same genotype as the mother (usually a carrier). Inference in the model aims at estimating the expected fractions of foetal- and maternal-specific DNA harbouring the HbS allele from which we predict the disease status of the foetus.

We assume that the read counts *r*_*i*_ and *s*_*i*_ follow Binomial distributions with parameters *θ*_*i*_ and *ϕ*_*i*_, the expected fractions of reads harbouring the HbS mutation in the mother and control, respectively:

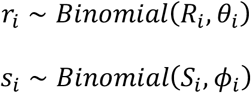

The fundamental intuition in the above model is to express *θ*_*i*_ as a mixture of maternal *ϕ*_*i*_ and foetal *ψ*_*i*_ fractions in proportions determined by the fraction *w* of foetal DNA in the mother’s blood, as follows:

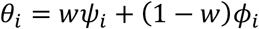

In the above mixture, the expected fraction *ϕ*_*i*_ is shared with the control and it can be estimated with relatively high confidence. We complete the model by imposing Beta priors (parametrized by mean and variance) on *ϕ*_*i*_ and *ψ*_*i*_, as shown below:

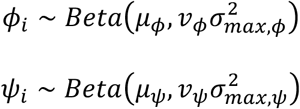

where *μ*_*ϕ*_ ~ *Uniform*(0, 1), *υ*_*ϕ*_ ~ *Uniform*(0, 1), and 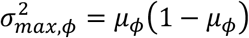. Analogous expressions hold for parameters *μ*_*ψ*_, *υ*_*ψ*_ and 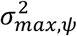. The above model was encoded in Stan and estimated using an adaptive Hamiltonian Monte Carlo algorithm with default parameters^14^.

Having estimated the foetal and maternal fractions of reads for each amplicon *i*, we need to decide whether the foetus is homozygous mutant (SS) and, therefore, the disease is present, or not (not SS). Assuming the mother is a carrier (i.e. she has genotype AS), we expect the fractions *ϕ*_*i*_ to be on average close to 50%. If the foetus is SS, then the foetal fractions *ψ*_*i*_ should be on average higher than *ϕ*_*i*_. We can quantify the difference between foetal and maternal expected read fractions by calculating the overlap between the above Beta distributions, as follows:

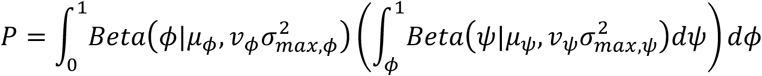

This measures the average probability that the foetal fraction is higher than the maternal fraction. Since the parameters of the above Beta distributions are random variables themselves, *P* is also a random variable with expectation 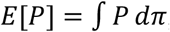, where *π* ≡ *π*(*μ*_*ϕ*_,*υ*_*ϕ*_,*μ*_*ψ*_,*υ*_*ψ*_) stands for the posterior distribution of its arguments. Given the output of the MCMC sampler, this expectation can be readily calculated as follows: a) for the *t*-th posterior sample of the parameter vector (*μ*_*ϕ*_,*υ*_*ϕ*_,*μ*_*ψ*_,*υ*_*ψ*_), sample one or more values of *ϕ* from its generative Beta distribution and, for each such value, calculate the inner integral in the above equation as a regularised incomplete beta function (this can be done efficiently using the *pbeta* routine in R), b) take the average of this integral over all samples of *ϕ*. This constitutes a posterior sample of *P*, corresponding to the *t*-th posterior sample of the parameter vector. c) Repeat until all *T* posterior parameter samples are exhausted. The required expectation can be calculated as an average over all *T* posterior samples of *P*.

Subsequently, we can say that the foetus has the disease (i.e. its genotype is SS), if the expectation of *P* exceeds a threshold *c*, i.e. *E*[*P*] > *c*. In order to find an optimal value of *c*, we utilise a training dataset with n=29 subjects and we identify the value of *c* in the range from 0 to 1 (in steps of 0.01), which maximizes the classification performance of the statistical model, as measured by the Matthews Correlation Coefficient (MCC). The variance of MCC was determined using 1000 bootstrap samples, each with size equal to the size of the training dataset (Figure 2D). The trained model was subsequently tested on a validation cohort with n=28 subjects.

